# *In host* mutational adaptation of *Mycobacterium tuberculosis* complex strains

**DOI:** 10.1101/2025.10.24.684346

**Authors:** Helen Zhang, Naomi Medina-Jaudes, Alicia Forcada-Nadal, Ewan M Harrison, Francesc Coll

## Abstract

**Background:** Tuberculosis (TB) is the second leading cause of death from an infectious disease worldwide. *Mycobacterium tuberculosis* complex (*MTBC*), the causative bacteria of TB, is becoming increasingly resistant to anti-TB drugs, resulting in poorer treatment outcomes. The mutations arising in *MTBC* strains during infection provide a record of bacterial diversification and adaptation, and indirect evidence on the selective pressures and conditions that *MTBC* encountered *in vivo*, such as those exerted by the immune system and antibiotic therapies.

**Methods:** We conducted a meta-analysis of *MTBC* sequenced strains with multiple clinical isolates sequenced from the same patient, from published studies and TB Portals. We applied stringent genomic QC criteria to keep only clonal isolates and applied robust genomic pipelines to identify mutations arising *de novo* during infection. A convergent evolution approach was applied to identify heavily mutated genes, operons, and candidate promoter across strains of multiple patients. We estimated the frequency of drug resistance acquisition during treatment for the subset of patients with available drug treatment data.

**Result:** Using 5,899 high-quality genomes from 1,056 TB patients after ensuring clonality of isolate genomes we identified limited within-host diversity was identified including 3,296 fixed mutations across 501 patients. A total of 21 genes, 25 operons and 27 promoter regions were statistically enriched by mutations compared to the rest of the genome, and additional loci with established or plausible adaptive roles approaching statistical significance. Significantly, this included multiple loci known to be involved in resistance to first-, second-, and last-line anti-TB drugs. Fluoroquinolone resistance was acquired more frequently during treatment than resistance to any other anti-TB drug. Previously reported candidate drug-resistance and -tolerance genes (*prpR*, *Rv2571c*, *fadD11*, *helY*, *ndhA*, *Rv0139*, *fadE5,* and *mce1* operon) were also identified. Genes encoding regulators (*phoR, whiB6* and *mycP_1_*) and effectors (*espK* and *eccE_1_*) of the virulence ESX-1 locus were frequently mutated *in host*.

**Conclusions:** Here we analysed a large dataset of *MTBC* within-host genetic diversity. We show that frequently mutated genes in *MTBC* during infection reveal known and biologically plausible *in host* adaptations, predominantly associated drug resistance, but also in genes involved in pathogenesis. The higher resistance acquisition rate observed for fluoroquinolones may have important clinical relevance. We reveal a list of candidate loci which will require mechanistic characterisation and whose impact on disease progression will need to be investigated.

## Introduction

The application of comparative, evolutionary and functional genomic approaches have accelerated the discovery of novel drug resistance (DR) loci in the genome of *M. tuberculosis complex* (*MTBC*), the causative pathogen of Tuberculosis (TB), beyond the canonical DR loci.^1^ Genome-wide association studies (GWAS) of drug-resistant and -sensitive strains have uncovered loci and genetic variants associated with DR^2–8^, many of which have been experimentally confirmed to mediate drug tolerance or resistance *in vitro*.^2,3,8^ Genome-wide evolutionary approaches, such as those based on metrics of selection^9,10^ or phylogenetic convergence^11,12^, have also been successful at revealing candidate loci relevant for host-pathogen interactions and DR.

Many previous studies have investigated *MTBC* within-host diversity and evolution using whole-genome sequencing (WGS), most often to distinguish relapse versus re-infection in cases of recurrent TB^13,14^, to quantify the degree of within-host genetic diversity^15,16^, and to study DR^17,18^ and heteroresistance evolution^19,20^ by inspecting mutations in known DR loci (reviewed here^21^). However, fewer studies have leveraged within-host diversity to study *in host* mutational adaptation of *MTBC* across the entire genome.^22–24^

Due to the strong evolutionary pressures exerted by prolonged anti-TB drug therapies, within-host evolution studies often identify genes involved in drug resistance or tolerance as heavily mutated or under parallel evolution. A previous study^24^ examined within-host diversity in *MTBC* strains from 200 patients with persistent or relapsed clonal infections and found seven known DR genes and three genes of unknown function with higher mutational density or under parallel evolution. Another study analysed a dataset of over 51,000 *MTBC* clinical isolates to identified unfixed (heterozygous) mutations as a surrogate for within-host diversity, to then identify genes repeatedly mutated across multiple patients.^22^ Known DR genes and transcriptional regulators were enriched among loci under positive selection, including a newly characterised transcriptional regulator with mutations that resulted in improved growth recovery after antibiotic exposure.

Among comparative and population-based studies, within-host evolution studies have been particularly successful at identifying mutational adaptations in bacteria.^24–26^ Because of the limited genetic diversity fixed in clonally evolving strains during infection, the power to detect signals of convergent evolution and positive selection is high when clonal genomes from the same strain and host are compared, and if large number of strains/hosts are considered. The mutations fixed in the genomes of clinical strains provide a record of bacterial diversification and adaptation *in host*, and indirect evidence of the selective pressures bacteria encounter *in vivo*. Because *MTBC* is an obligate human pathogen^27^, all selective forces acting on *MTBC* are expected to originate within the host, including those exerted by antibiotic therapies, nutritional limitations of the host and the immune system.

Here we analyzed a dataset of 5,800 *MTBC* genomes from more than 1,000 TB patients sourced from published studies, and applied a genome-wide mutation enrichment approach^26^ to identify mutational adaptations. Overall, we identified 21 genes, 25 operons and 27 promoter regions in the *MTBC* genome statistically enriched by mutations, and more loci with established or plausible adaptive roles approaching statistical significance. In addition to many canonical DR loci, we also found candidate DR loci (*prpR/Rv1129c*, *Rv2571c*, *fadD11/Rv1550*, *helY/Rv2092c*, *ndhA/Rv0392c*, *Rv0139*, *fadE5/Rv0244c,* and *mce1* operon) identified elsewhere by independent GWAS and functional genomics studies. Putative adaptive variants in loci regulating ESX-1 secretion at both transcriptional (*phoR, whiB6*) and post-transcriptional levels (*mycP1*), as well as in effector ESX-1 encoded proteins (*espK* and *eccE_1_*) were identified. Notably, among all DR genes, *gyrA* was by far the most mutated locus, attributable to a higher *de novo* acquisition rate of fluoroquinolone resistance.

## Results

### Characteristics of the compiled dataset of M. tuberculosis complex genomes

A total of 7,011 unique *MTBC* isolate genomes were identified from the NCBI in April 2022 from TB Portals (n=856)^34^ and a total of 34 different studies (n=6,155)^13,16,17,19,20,28–33^ (studies with >20 patients cited, see Supplementary Table 1 for all) -which originated from 1,409 different TB patients who had multiple isolates sequenced (median of 2 isolates per host, 2-3 IQR). An additional 5,518 isolates from other patients in the same studies, who had a single *MTBC* genome sequenced per host, were additionally considered as contextual isolates (Supplementary Data 1 and Supplementary Table 1). Stringent genomic QC was applied to discard cross-contamination with non-*MTBC* DNA, mixed infections of multiple *MTBC* strains, and to ensure clonality of isolates from the same host (see Methods and Supplementary Figure 1). This stringent QC was needed to minimize the rates of calling false-positive variants,^24^ a problem amplified when comparing highly related clonal genomes from the same strain. After genomic QC, 5,882 *MTBC* genomes (81.8%) from 1,044 hosts (70.8%) were kept (Figure 1A), plus an additional 5,417 contextual isolates.

**Figure 1.**
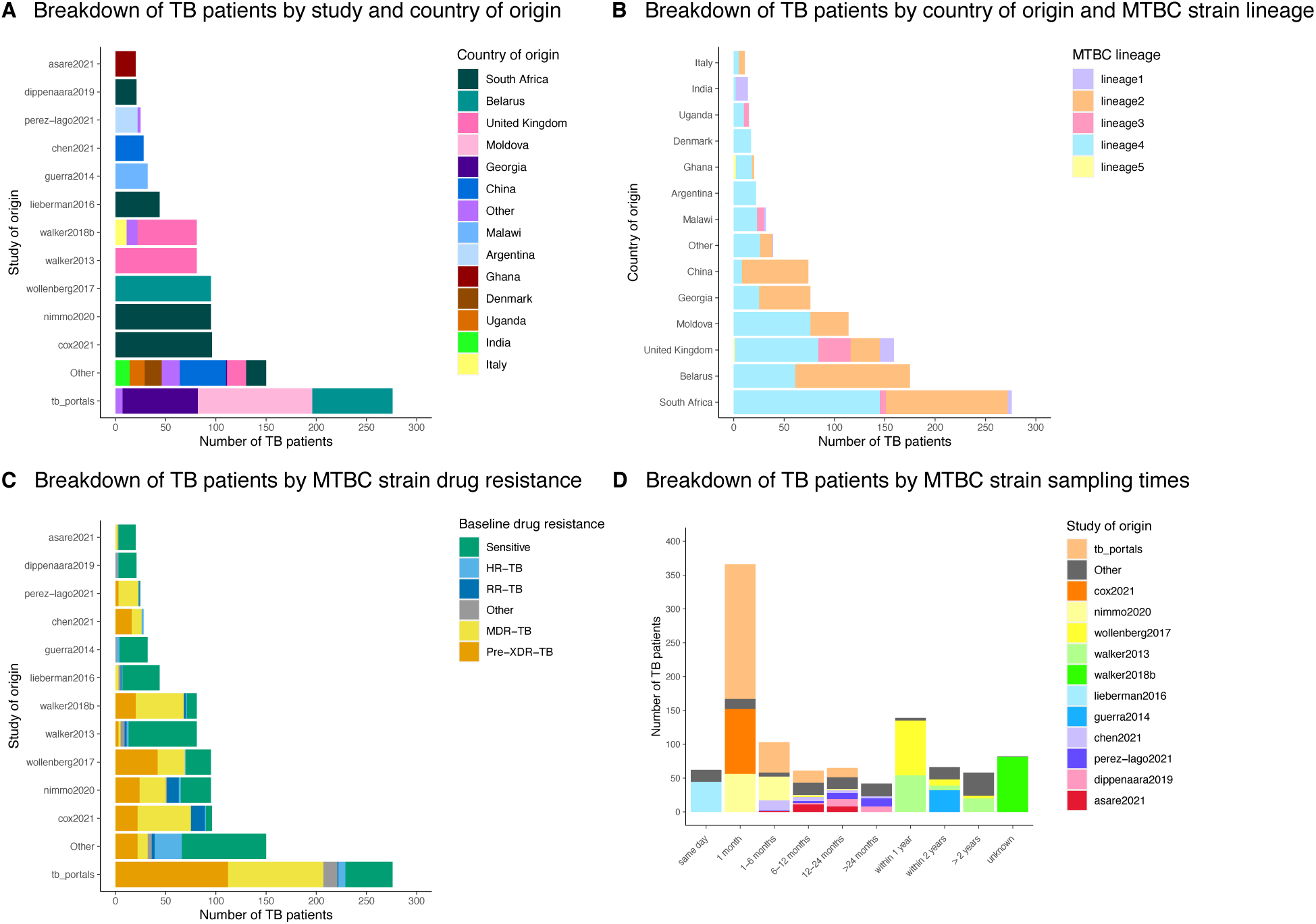
Characteristics of compiled MTBC strain dataset used in this study. **A** Breakdown of TB patients by study and country of origin. The share of TB patients by study is color-coded by country of origin. Studies contributing less than 20 TB patients and countries contributing less than 10 patients are labelled as “Other”. **B** Breakdown of TB patients by country of origin and MTBC strain. The share of TB patients by country of origin is color-coded by MTBC lineage, as determined from genome sequences by TB-Profiler. **C.** Breakdown of TB patients by the baseline DR of their MTBC strain. The share of TB patients by study is color-coded by baseline genotypic DR, as determined from genome sequences by TB-Profiler. Baseline DR is defined considering all isolates genomes from the same strain and host, and assuming the least DR patter is that of the original (baseline) infecting strain. **D.** Breakdown of TB patients by MTBC sampling times. The maximum time distances between sample collection times of all isolates from the same strain and host are shown in the x-axis. The share of TB patients by collection time distances are color-coded by study.

In terms of geographical representation, *MTBC* strains originated from 27 different countries (Figure 1A), mostly sampled from Europe (578, 55.4%) and Africa (343, 32.9%). In Europe, most TB patients were sampled from Belarus (175, 16.8%), the United Kingdom (159, 15.2%), Moldova (114, 10.9%), and Georgia (76, 7.3%); and in Africa mostly from South Africa (276, 26.4%). As expected, most *MTBC* strains belonged to lineage 4 (519 out of 1,044, 49.7%) and 2 (439, 42.0%), while the rest were assigned to lineage 3 (50, 4.8%), 1 (33, 3.2%) and 5 (3, 0.3%) (Figure 1B). Most patients had isolates grown from pulmonary specimens (578, 55.4%), most often from sputum specimens only (526, 50.4%), followed by both sputum and bronchoalveolar lavages (16, 1.5%). Only 1 patient had extra-pulmonary specimens only, and 56 had both pulmonary and extra-pulmonary specimens. The anatomical origin of samples was not reported for 408 patients (39.1%).

In terms of baseline genotypic DR, there was an even and sizable representation of pan-susceptible (370, 35.4%), MDR (296, 28.4%), and pre-XDR strains (264, 25.3%). Among the other DR patterns (114, 10.9%), strains were mostly mono resistant to either isoniazid (49, 4.7%) or rifampicin (36, 3.4%) (Figure 1C). In terms of timing of collection between samples, these were available for most patients (699, 67%), of which 62 (5.9%) had samples collected on the same day, 366 (35.1%) within one month, 164 (15.7%) between one to twelve months, and 107 (10.3%) within more than one year apart (Figure 1D). For patients with unknown sampling dates (345, 33%), 139 (13.3%) patients had their samples collected within one year, 66 (6.3%) within two years, and the rest (58, 5.5%) within more than two years apart. The remaining 82 patients (7.9%) had no available information on sampling collection dates.

### Genome-wide mutation enrichment analysis identifies robust signals of in host adaptation

Using clonal isolates sampled from the same host, we quantified the number of protein-altering mutations (missense, nonsense, and frame-shift mutations) within each protein coding sequence (CDS) that arose *de novo* during in *MTBC* strains during TB infections. In RNA coding genes and intergenic regions, we considered all mutations regardless of their impact annotation. Limited within-host diversity was identified including 3,296 unique fixed mutations across 503 patients (out of 1,044), with a median of 2 fixed mutations per strain (1-5 IQR). To identify loci in the *MTBC* genome exhibiting evidence of adaptation, we applied a genome-wide mutation enrichment approach that identifies signal of parallel and convergent evolution across strains of multiple individuals, as previously implemented.^26^ We excluded repetitive regions and regions difficult to genotype using Illumina short-read data to minimize the rates of calling false-positive variants and inflating mutation counts. Out of the 4,173 genes annotated in the H37Rv reference genome (including 142 RNA and 4,031 protein coding genes), a total of 21 genes were found to be statistically enriched by mutations compared to the rest of the genome (Figure 2A). Not surprisingly, a large proportion of these genes (16/21, 76%) are known to be involved in DR^1^ and most likely represent adaptations of *MTBC* strains to survive anti-TB therapies, demonstrating the robustness of our approach in detecting expected adaptive mutations.

**Figure 2.**
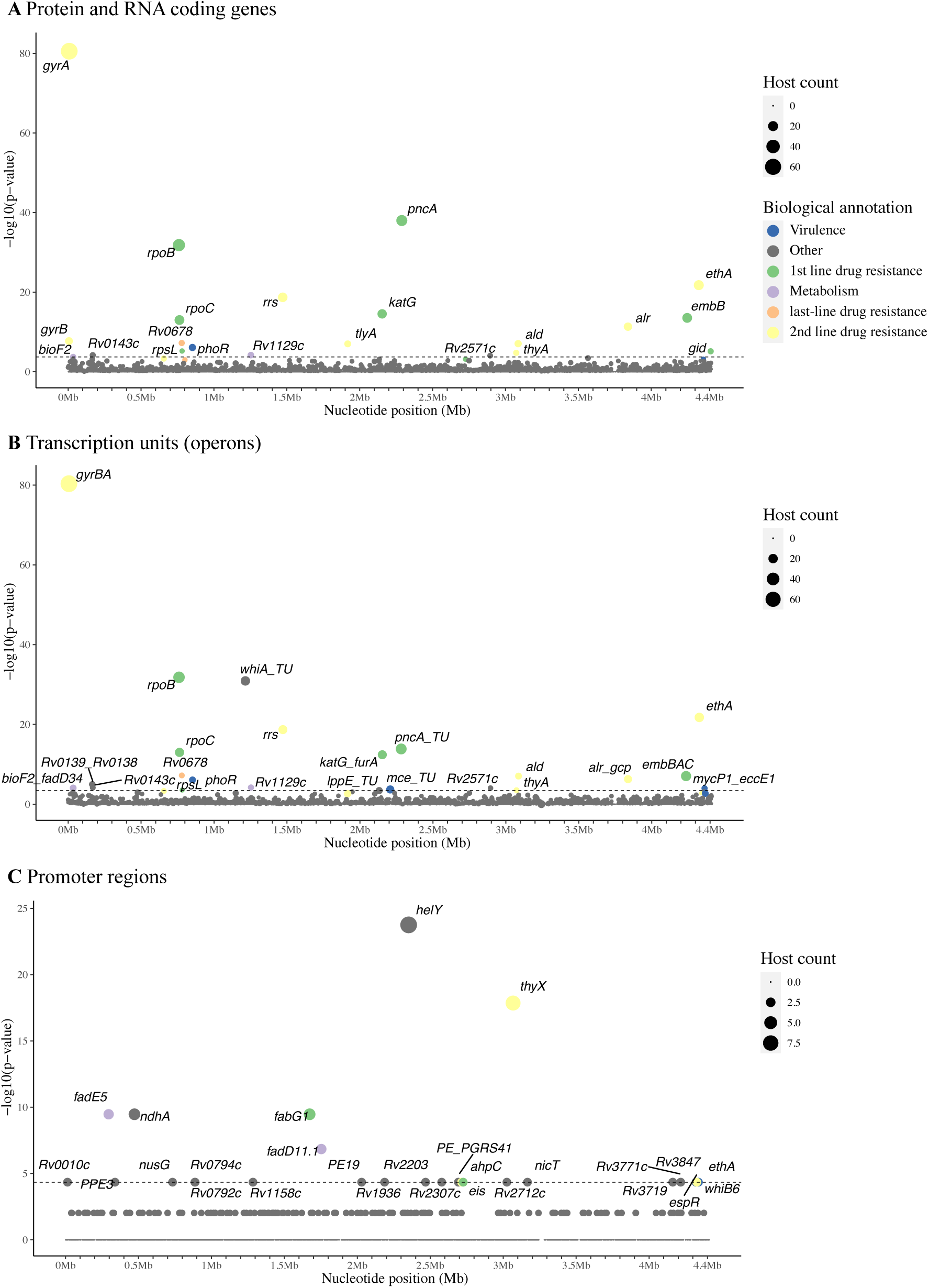
Loci enriched with mutations across multiple MTBC strains. **A** Protein and RNA coding genes, (**B**) transcriptional units (operons), and (**C**) promoter regions (100 bp upstream of annotated operons) enriched for mutations across the MTBC strains of multiple TB patients. Each circle denotes a single locus, whose size is proportional to the number of hosts mutations arose independently from. Loci are placed at the x-axis based on their chromosome coordinates. Loci are colour-coded by biological functions, particularly if they are known to be involved in DR or virulence. The y-axis shows the uncorrected *p*-value resulting from a single-tailed Poisson test comparing the density of mutations per functional unit against the expected number of mutations, obtained from multiplying the genome-wide mutation count per bp by the gene length (see Methods). The dotted horizontal line represents the genome-wide statistical significance threshold. Mutations identified in a collection of 5,882 *MTBC* isolate genomes from 1,044 individuals (see Supplementary Table 1).

To broaden the search for signals of convergent evolution, we counted mutations among all genes annotated on the same transcription unit (operon). Out of the 2,573 operons in the reference genome, 23 reached statistical significance for an excess of mutations (Figure 2B), most often containing genes already identified on their own (Figure 2A). Five additional operons contained genes that did not reach statistical significance independently, including *Rv0138-Rv0139*, *PE10 (Rv1089)*, *lppE (Rv1881c)* – *fbpB (Rv1886c)*, *yrbE3A* (*Rv1964)* -*Rv1975* (*mce1* operon) and *mycP1* (*Rv3883c*) *-eccE1* (*Rv3882c*). When considering putative promoter regions, 27 regions reached statistical significance (i.e., with at least two strains acquiring mutations independently), including many promoter regions known to be involved in DR (*ahpC*, *eis*, *thyX*, *thyA* and *fabG1-inhA*) (Supplementary Table 2).

### Signals of in host mutational adaptation reveal well-known drug resistance loci

As expected, many of the loci with signals of *in host* mutational adaptation were in canonical DR loci. Among these we identified loci linked to resistance to first-line drugs (*katG*, *ahpC* promoter, *inhA* promoter, *rpoB*, *rpoC, pncA* and *embB*), second-line drugs (*gid, rpsL, rrs, eis* promoter, *tlyA, gyrA*, *gyrB*, *ethA* and *ethA* promoter) and last-line drugs (*alr*, *ald, Rv0678, thyA* and *thyX* promoter) (See Supplementary Table 2 for corresponding drugs). Other well-characterised DR loci were found among the top mutated genes approaching statistical significance, including genes involved in resistance to isoniazid (*inhA, ahpC, Rv2752c* and *ndh*), ethionamide (*Rv0565c* and *inhA*), and linezolid (*rplC*). Notably, among all DR genes, *gyrA* was by far the most mutated locus. The high number and diversity of known resistance mechanisms identified is reflective of the size of the analysed dataset (1,044 TB patients), and the diversity of strains’ baseline DR profiles (including pan-susceptible (370, 35.4%), MDR (296, 28.4%), and pre-XDR strains (264, 25.3%)). Next, we compared individual mutations in these DR loci against those in the WHO catalogue of DR mutations^1^, except for loci associated with resistance to cycloserine (*ald* and *alr*) and para-aminosalicylic acid (*thyA* and *thyX* promoter) which are not included in the catalogue. We found a total of 323 independently acquired mutations in 164 different strains (218 unique variants across all strains) in 31 DR loci (Supplementary Data 2). Out of these 218 unique variants, 35 (16%) are not listed in the WHO catalogue. Out of the 183 (83.9%) listed, 93 (42.7%) are reported to be associated with resistance, 7 (3.2%) not associated, and 83 (38.1%) associated with uncertain significance.

### In host mutational adaptation reveals less characterized and candidate drug resistance loci

The genes *prpR* (*Rv1129c*), *Rv0143c*, *Rv2571c* and *bioF2 (Rv0032)* were statistically enriched by mutations *in host*, as many canonical DR genes were (Figure 2A). Mutations in *prpR*, a transcription factor that regulates propionate metabolism, have been reported to be prevalent in clinical *MTBC* strains and to confer drug tolerance to multiple drug classes.^2^ In a GWAS of MDR-TB clinical isolates from Peru, *Rv2571c* was identified to be strongly associated with ethambutol resistance^5^ and mutations in *bioF2* (*Rv0032*) have been hypothesized to be implicated in increasing levels of streptomycin resistance.^35^ No role in DR has been proposed for *Rv0143c*, a gene of unknown function that possibly encodes a transmembrane chloride channel transporter.

When considering candidate promoter regions, the intergenic region upstream of *helY* (*Rv2092c*), which encodes for the ATP-dependent DNA helicase HelY, was the most mutated promoter region (Figure 2C). *helY* promoter mutations may well be involved in DR as its coding region (*Rv2092c*) was previously found to be associated with isoniazid resistance in a GWAS.^2^ Related to previous observations that mutations in the coding region of *ndhA* (*Rv0392c*) are phylogenetically associated with resistance, and that *ndhA* transposon mutants exhibit increased isoniazid resistance^36^, we found an enrichment of mutations in the *ndhA* promoter region, in addition to the *ndh* coding region (approaching statistical significance), both sharing high amino acid similarity and encoding NADH dehydrogenases. We additionally identified the promoter regions of *fadE5* (*Rv0244c*) and *fadD11-plsB1* (*Rv1550-Rv1551*) with evidence of convergent evolution (Figure 2C). Overexpression of *fadE5* in *M. smegmatis*, encoding an acyl-CoA dehydrogenase, has been shown to increase resistance to ethambutol and streptomycin^37^; while knockout mutants of *fadE5* in *M. tuberculosis* exhibit reduced tolerance to rifampicin.^38^ A recent GWAS found that large deletions and loss-of-function mutations in *fadD11* are associated with streptomycin resistance.^4^

When considering transcriptional units, the *mce1* operon (*Rv1964* to *Rv1975*) was among the top mutated operons (Supplementary Data 3), embedded within operons containing canonical DR genes (Figure 2B). Transposon mutagenesis has shown that knockouts of *mce1* operon genes display increased isoniazid resistance^36^, consistent with their role in cell wall lipid transport and homeostasis.^39^

### Loci encoding regulators of virulence-associated ESX-1 locus are often mutated in host

The most frequently mutated gene *in host* beyond canonical DR genes was *phoR*, the gene encoding the sensor kinase of the *phoPR* two component system which controls the secretion of ESAT-6, considered one of the major virulence factors of *M. tuberculosis*. Consistent with this finding, previous genomic studies have found strong evidence of positive selection in *phoR*^10,40^, particularly in the sensor domain. We identified PhoR mutations across the full-length protein (Figure 3A) including a substitution on the sensor domain (Arg80Met); another in the HAMP domain (Ala221Val), responsible for signal transduction from the transmembrane to cytoplasmic catalytic regions; and three mutations clustered within the dimerization-phosphorylation domain (Pro264Leu, Tyr275Asp and Ala279Val), of which Pro264Leu could impair the conformational flexibility essential for transmitting conformational changes from the sensor domain through the transmembrane and cytoplasmic helices to regulate kinase activity.^41^ Finally, three additional mutations (Leu324Arg, Arg335Trp and Gly350Cys) were found in the ATPase domain which could interfere with ATP binding, domain stability, or interdomain coupling required for efficient kinase activity.

**Figure 3.**
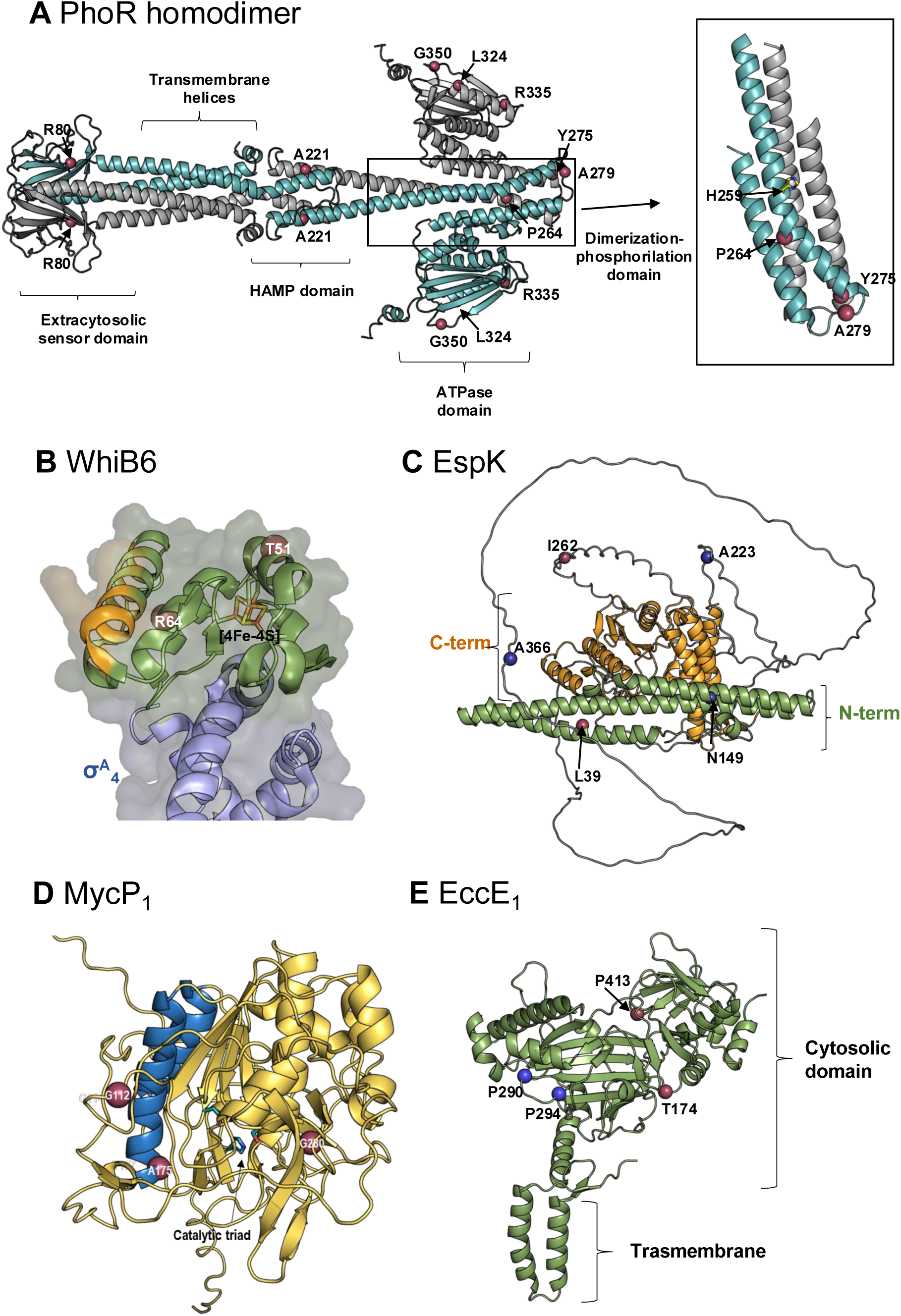
Localization of the protein structure of mutations in ESX-1-associated proteins. **A** Structural model of the *M. tuberculosis* PhoR homodimer predicted by AlphaFold. One subunit is shown in green and the other in grey. Mutations identified in this study are highlighted as red spheres. The inset shows the dimerization–phosphorylation domain from the crystal structure (PDB 5UKV), with mutation positions indicated in a single subunit for clarity. The active-site His259 residue involved in autophosphorylation is shown as sticks. **B** Structure of the *M. tuberculosis* WhiB6 (green)–σA4 (blue) complex (PDB 8DV5). The identified mutations are shown as red spheres. The [4Fe–4S] cluster and its coordinating cysteine residues are represented as sticks, and residues predicted to interact with DNA are highlighted in yellow. **C** AlphaFold model of *M. tuberculosis* EspK. The N-terminal domain is shown in green, the C-terminal domain in yellow, and the long flexible linker in grey. Point mutations are represented as red spheres and frameshift mutations as blue spheres. **D** AlphaFold model of *M. tuberculosis* MycP1. Mutated residues are shown as red spheres. The catalytic triad is represented as green sticks, and the helices involved in substrate binding are highlighted in blue. The transmembrane helix is omitted for clarity, as no mutations were located in this region. **E** AlphaFold model of *M. tuberculosis* EccE1. Residues bearing point mutations are shown as red spheres, and those with frameshift mutations as blue spheres.

Functionally related to mutations in PhoR, we found an enrichment of protein-altering mutations in WhiB6 (*Rv3862c*) and in the ESX-1 secretion-associated protein EspK (*Rv3879c*), both approaching statistical significance (Supplementary Data 3). *whiB6* is located adjacent to the ESX-1 locus and is another known ESX-1 transcriptional regulator. The expression of *whiB6* is, in turn, positively regulated by PhoP which binds directly to its promoter^42,43^; and EspK is known to be transcriptionally regulated by WhiB6. Figure 3B shows that WhiB6 substitutions (Thr51Pro, detected twice, and Arg64His) are not located within the regions predicted to interact with DNA^44^ or with the RNA polymerase α-subunit^45^ but within α-helices that contain conserved cysteines coordinating the [4Fe-4S] cluster. Interestingly, a recent study^46^ found 13 independent WhiB6 variants linked to decreases in PE35-PPE68-esxBA operon expression, including substitutions in WhiB6 close to the one detected here (Cys53Gly, Asp46Ala). EspK is an auxiliary protein that recruits ESX-1 substrate EspB through its C-terminal domain, while its N-terminal domain interacts with the core secretion component EccCb_1_ to guide EspB towards the secretion pore.^47,48^ We identified five mutations in EspK (Figure 3C): two missense variants (Leu39Trp and Ile262Thr) and three frameshifts (Gln149fs, Ala223fs, and Ala366fs), the latter predicted to produce truncated and non-functional EspK that would compromise EspB secretion.

When considering transcriptional units, the operon encoding MycP_1_ (*Rv3883c*) and EccE_1_ (*Rv3882c*) (TU185E-7171) was statistically enriched by protein-altering mutations. MycP_1_ is a serine protease that regulates substrates of ESX-1 like EspB post-transcriptionally^49^, and EccE_1_ an effector membrane component of ESX-1 required for secretion of ESX-1 substrates.^50^ In MycP_1_ we identified three missense mutations (Gly112Asp, Ala175Gly, and Gly280Ser), the first two within or adjacent to the α-helices forming the substrate-binding pocket but distant from the catalytic triad (Figure 3D), thus not expected to directly affect proteolytic activity but might influence substrate recognition or enzyme dynamics. In EccE_1_ we identified four mutations: two missense substitutions (Thr174Pro and Pro413Thr) in surface-exposed loops of the cytosolic domain (Figure 3E), where they could affect local flexibility or protein-protein interactions; and two frameshift variants (Pro290fs and Pro294fs) predicted to generate truncated proteins.

Taken together our observations point to an important role of adaptive mutations in ESX-1 loci, in distinct yet interconnected layers of ESX-1 regulation: PhoR and WhiB6 which control transcriptional activation, EccE_1_ and MycP_1_ which form part of the core secretion machinery, and EspK which operates as an accessory adaptor linking substrates such as EspB to the translocon. Mutations identified in these genes could therefore influence ESX-1 activity at multiple levels, from gene regulation to substrate recognition and secretion.

### Fluoroquinolone resistance was acquired more frequently than resistance to any other drug

We observed that *gyrA* was by far the most frequently mutated gene (Figure 2A), therefore we investigated whether a more frequent use of this drug in treatment or a higher rate of resistance acquisition in the dataset analysed could explain this observation. Drug treatment data was available and could be extracted for 550 (52%) of the 1,066 strains analysed, being mostly available from TB Portals (51%) (Supplementary Table 3). Most strains were sampled from Belarus (31.1%), Moldova (20.7%), South Africa (19.3%) and Georgia (13.6%), and had a distribution of baseline genotypic DR that mirrored that of the overall dataset (31.6% pan-susceptible, 28.7% MDR and 23.5% XDR) (Supplementary Figure 2). Strains were treated with a range of anti-TB drugs, most often with pyrazinamide (n=438, 79.6%), fluoroquinolones (n=352, 64.0%), and ethambutol (n=298, 54.2%) (Table 1). Of these 550 strains, 83 (15%) had at least one fixed *de novo* mutation in a DR gene. Fluoroquinolone resistance appeared to be acquired more often than to other drugs: of the 352 strains treated with fluoroquinolones 46 acquired a fixed DR mutation in *gyrA* or *gyrB* (13.1%, 95% CI 9.9 –17.0%, see Table 1). This proportion was higher than that observed for other drugs, even for first-line drugs that many strains were treated with such as isoniazid (3.5%, 95% CI: 1.7 – 6.8%), rifampicin (1.6%, 95% CI: 0.5 – 5.0%), ethambutol (2.7%, 95% CI: 1.3 – 5.3%) and pyrazinamide (3.7%, 95% CI: 2.2 – 5.9%). The acquisition rate for fluoroquinolone resistance was higher among MDR strains (17.9%, 95% CI: 12.5 - 25.1%).

**Table 1.**
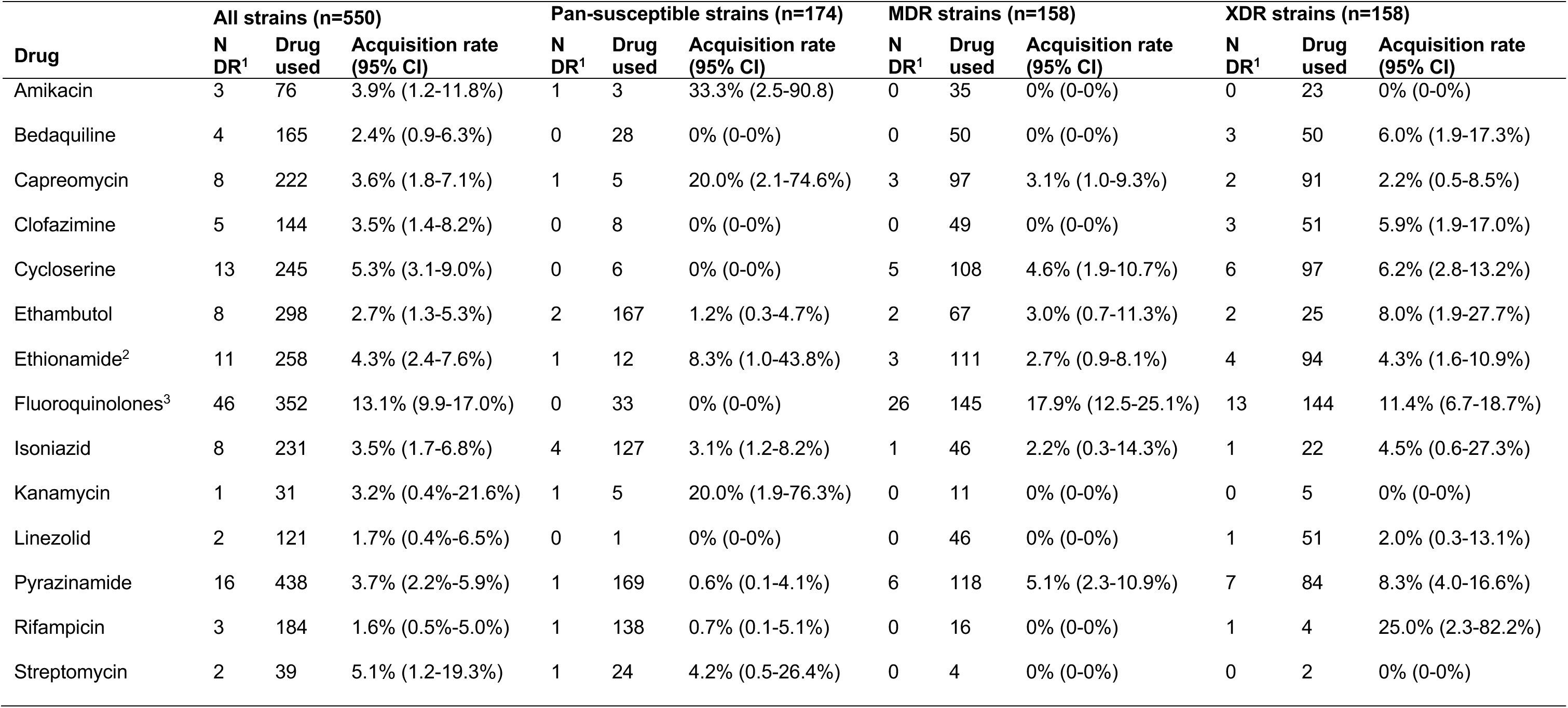
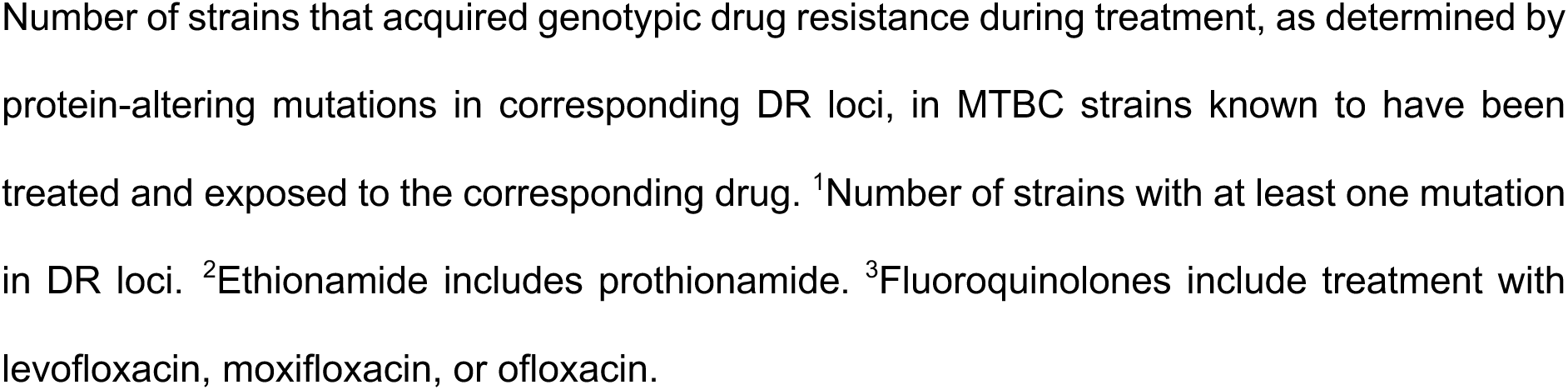
Frequency of genotypic drug resistance acquisition in MTBC strains.

## Discussion

Here we conducted a meta-analysis of published collections of multiple *MTBC* isolates sequenced from the same TB patient to capture within-host genetic diversity and identify signals of *in host* adaptation. After stringent genomic quality control, we kept over 5,800 *MTBC* genomes from more than 1,000 TB patients, sourced from TB-Portals and published studies, and of diverse phylogenetic and geographical origins. Using clonal isolates from the same host, we identified mutations that arose *de novo* in *MTBC* strains during infection. We next applied a genome-wide mutational enrichment approach to identify loci in the *MTBC* genome under parallel and convergent evolution that could represent adaptations during TB infection, providing indirect evidence of the stressors and evolutionary pressures that *MTBC* strains encountered *in host*. Overall, we identified 21 genes, 25 operons and 27 candidate promoter regions in the *MTBC* genome statistically enriched by mutations, and more loci with an established or plausible adaptive role approaching statistical significance.

The identification of many canonical DR genes demonstrated the robustness of our approach in detecting expected mutational adaptations. These confirmatory results give credibility to other putative adaptations identified in our study, mostly in ESX-1 loci and candidate DR loci. Among the latter, our results support the adaptive role of mutations in *prpR/Rv1129c* (which mediates tolerance to multiple drug classes)^2^, *Rv2571c* (previously associated with ethambutol resistance)^5^, *fadD11/Rv1550* (associated with streptomycin resistance)^4^ and *helY/Rv2092c* (reported to be associated with isoniazid resistance)^2^, all of them identified in GWAS of DR in clinical strains. However, caution should be taken when interpreting DR-locus associations in GWAS studies, as co-resistance patterns often observed in clinical strains can complicate these interpretations and thus associated DR loci could well be conferring resistance to another drug or multiple drugs.

Our results also support the findings of recent functional genomics studies of DR in *MTCB*. We found an enrichment of mutations in the promoter region of *ndhA/Rv0392c* (encoding an NADH dehydrogenase) and in the coding region of *Rv0139* (encoding a possible oxidoreductase), both linked to isoniazid resistance by transposon mutagenesis^36^, and consistent with the role of redox metabolism in isoniazid resistance.^51^ This same study^36^ identified transposon mutants in the *mce1* operon exhibiting increased isoniazid resistance, in line with *mce1* operon found among to most mutated operons in our study, and consistent with the role of cell wall lipid transport and homeostasis in isoniazid resistance.^39^ Still, considering that Mce systems are key virulence factors in *MTBC*, required for organized granuloma formation and persistence^52^, *mce1* operon variants could well be modulating *MTBC* pathogenesis too. We also identified the promoter region of *fadE5* (*Rv0244c*), whose overexpression increases resistance to ethambutol and streptomycin in *M. smegmatis*.^37^ All in all, our results show that within-host evolutionary studies can complement GWAS and transposon mutagenesis approaches in identifying new DR candidate loci.

Interestingly, we found that *gyrA* was by far the most mutated gene *in host* and that this was attributable to a higher acquisition rate of resistance to FLQs rather than just a more common exposure to this drug in the dataset analysed. Longitudinal sampling of *MTBC* have also showed that *de novo* acquisition of resistance is particularly high for fluoroquinolones.^17^ Similar rates of acquired FLQ resistance have been reported in studies of MDR TB (12.1% and 9.1%)^53,54^, although these studies reported higher acquired resistance rates to second-line injectable drugs (7.8% and 9.8%) than the ones detected here. The higher rate of *gyrA* mutations could be in part attributable to the common empirical use of FLQs to treat other community infections, as it has been shown that such prior exposure entails a higher risk of developing FLQ-resistant TB.^55^ Alternatively, higher rates of heteroresistance to FLQs^2,20^ may also explain their apparent higher resistance acquisition rate measured here.

Beyond resistance to anti-TB drugs, our results point to an important role of adaptive mutations in regulators and effectors of the ESX-1 secretion system, most often in *phoR* but also in other transcriptional (*whiB6*) and post-translational (*mycP_1_*) regulators and effectors (*espK* and *eccE_1_*) of this system. In this work, several mutations were identified in the *phoR* and *whiB6* genes which could modulate ESX-1 substrate secretion and directly impact host-pathogen interactions. Previous population and evolutionary studies have identified *phoR* as a target of positive selection^10,40^, and clinical strains of *M. abscessus* with prevalent PhoR variants have been reported and linked to an improved survival of strains within the host.^56^ ESX-1 substrates are not just key virulence factors but also highly immunogenic antigens.^57^ Therefore, MTBC must tightly regulate the activity of ESX-1 to balance virulence and immune recognition and ensure the maintenance of successful infections. Whether *phoR* mutations are a case of directional or balancing selection remains to be determined. *MTBC* strains could be evolving to enhance or attenuate their virulence *in host* (directional selection). Indeed, a recent study^46^ found repeated evolution of WhiB6 variants linked to decreased expression of virulence effectors secreted by ESX-1, expected to lead to attenuated virulence. The identification in our study of loss-of-function mutations in EspK and EccE_1_, an ESX-1 auxiliary and effector proteins, respectively, both required for secretion of ESX-1 substrates, also points towards an adaptation towards lower virulence. If balancing selection is at play, then genetic variants in ESX-1 regulators may allow *MTBC* cell populations to diversify *in host* and create subpopulations of cells that are preadapted to different conditions or immunological contexts. The presence of spatially separated bacterial subpopulations in different lung cavities with different immunological contexts could also lead to variation in ESX-1 regulators. The recent identification of common phase variants upstream of other ESX-1 regulators points to the importance of maintaining heterogeneity in ESX-1 expression *in host*.

Our study has several limitations. First, most *MTBC* strains had only two isolates sequenced per host available from published studies, limiting the amount of genetic diversity captured from clinical specimens. Sequencing directly from sputum^58^ in large cohorts of TB patients would provide more realistic estimates of the prevalence and acquisition rate of the adaptations identified here. Third, we only considered fixed genetic variants and excluded heterozygous variants, which can represent low abundant adaptive mutations on a trajectory of fixation^24^, fixed at certain anatomical sites^16^, or in balancing selection. Finally, while drug treatment data was available for a subset of patients, this only included which drugs were used and no details on the duration of exposure or changes of drugs administered during the course of treatment, which may have impacted our estimations of drug resistance acquisition.

Adaptation of clinical *MTBC* strains during infection has been the focus of multiple previous studies, which demonstrated the suitability of *in host* evolutionary approaches to discover new mechanisms of genetic adaptation, most often related to drug resistance. Here we conducted a meta-analysis of 5,882 *MTBC* genomes from 1,044 TB patients which revealed known and additional mutational hotspots in the *MTBC* genome, in loci of established and plausible adaptive roles during infection. The less characterised loci now warrant experimental validation to confirm the environmental stressors that selected adaptive mutations, the underlying molecular mechanisms mediating adaptation and survival during infection, and their impact on disease progression.

## Methods

### Data sources

We aimed to identify published collections of multiple *M. tuberculosis* complex (MTBC) isolates sequenced from the same TB patient, whether collected longitudinally or from the same specimen (same day). The NCBI Short Read Archive (SRA) was queried on April 2022 to identify BioProjects that met the following criteria: contained MTBC genomic sequences, could be linked to a published study, included genomes of clinical origin, multiple isolates per host were sequenced, and host IDs were available and could be linked to isolate IDs. In addition, MTBC genomes were obtained from TB Portals^34^ (https://tbportals.niaid.nih.gov/) to keep the subset of genomes from hosts with multiple isolates sequenced per host. In addition, we extracted the genomes and metadata of MTBC isolates from other patients in the same studies who had a single MTBC genome sequenced per host as contextual isolates. Isolate IDs (run accession, BioSample accession, sample name) and basic metadata (patient ID, isolation source, collection date and geographical origin) were extracted from published studies, the NCBI BioSample and TB Portals.

### Genomic analyses applied to all M. tuberculosis complex isolate genomes

The taxonomic classifier Kraken v2.1.2^59^ was used with default parameters and database (*kraken2-build --standard*) from raw fastq files to identify potential contamination with non-target DNA. TB-Profiler v4.2.0^60^ (https://github.com/jodyphelan/TBProfiler) was used to assign lineage and sub-lineages to MTBC genomes and identify drug-resistance mutations. Snippy v4.6.0 (https://github.com/tseemann/snippy) was used to map Illumina short reads to an ancestral version of the *M. tuberculosis* H37Rv reference genome^61,62^ and to call genetic variants (i.e., SNPs and short indels). Only functional mutations (i.e., any type of mutation other than synonymous amino acid changes) within protein coding sequences were included and all others within RNA-coding and intergenic regions. Illumina paired-end reads were assembled using a computational pipeline based on SPAdes assembler^63^ v3.15.3 and the improve_assembly pipeline (https://github.com/francesccoll/assembly_pipeline) designed to improve draft assemblies by scaffolding and gap filling. Illumina single-end reads were assembled using SKESA 2.4.0.^64^ The quality of all assemblies was evaluated using QUAST v5.0.1.^65^

### Genomic QC metrics and thresholds

Given the diversity of studies genomic data was identified from, we assessed a variety of genomic QC metrics and applied stringent QC thresholds to rule out cross-contamination with non-MTBC DNA and cases of mixed infections, and to ensure an overall high-quality of sequence data across studies. For defining genome QC thresholds, we considered all MTBC genomes available from the identified BioProjects, regardless of whether linked isolate-level metadata could be extracted or not. From Kraken2 reports the proportion of MTBC reads and that of the most abundant contaminating species was extracted. From TB-Profiler output, the presence of more than one *M. tuberculosis* sub-lineage (at the lowest sub-lineage level) was used as evidence of mixed infections. From Snippy derived *bam* and *vcf* files, we extracted the number of heterozygous sites and average sequencing depth. From QUAST reports we considered the total number of contigs, the total number of bases in the assembly and N50. We plotted the values of each of these metrics for all genomes to identify clear outliers and thresholds separating them form the rest of genomes (Supplementary Figure 1). Low-quality genomes were excluded from further analysis by applying the following thresholds: Kraken proportion of *M. tuberculosis* complex reads < 90%, average sequencing depth < 20%, percentage of missing call > 10%, heterozygous sites ratio > 0.002, number of contigs > 1,000, N50 < 150,000 bp and total assembly length < 4,0 Mb or > 5,5 Mb.

### Genomic analyses applied to isolates of the same host

To avoid comparing the genomes of divergent strains from the same individual (i.e., mixed infections), only clonal isolates were kept for further analyses. Clonality was ruled out if isolates belonged to different TB-Profiler sub-lineages (at the lowest sub-lineage level) or to the same sub-lineage separated by more than 10 SNPs, a stringent genetic relatedness distance equivalent to the previously established cut-off for *M. tuberculosis* cross-transmission.^30^ We applied a previously developed pipeline^26^ to identify *de novo* genetic variants originating during infection from multiple isolates sequenced from the same host. In short, the nucleotide sequence of the most recent common ancestor (MRCA) of all isolates from the same host was reconstructed with PastML v1.9.20^66^ from within-host phylogenies built with *RAxML* v8.2.8^67^, rooted on a closely related contextual isolate sampled from a different host (outgroup). This pipeline was implemented in four python scripts (identify_host_ancestral_isolate.step1.py to identify_host_ancestral_isolate.step4.py) available at https://github.com/francesccoll/mtbc-adaptive-mutations/. Variants in repetitive regions, detected by running Blastn v2.8.1+ on the reference genome against itself, and variants in regions of low complexity, as detected by *dustmasker*^68^ v1.0.0 using default settings, were also filtered out. We additionally masked regions defined as repetitive regions elsewhere and low confidence regions that are difficult to genotype using Illumina short reads.^69^ Overall, these masked regions accounted for a total of 735,968 bp (16.7% of the H37Rv reference genome) and were excluded from further analysis. The final set of high-quality variants were annotated using *SnpEff* v4.3^70^ in the H37Rv reference genome.

### Genome-wide mutation enrichment analysis

A convergent evolution approach was applied to identify heavily mutated loci across multiple patients, to identify putative adaptive genetic changes as previously described. Here, only fixed mutations called by Snippy were considered. We counted protein-altering mutations (i.e., those annotated as having HIGH or MODERATE annotation impact by *SnpEff*) within protein coding sequence and operons, and all mutations within RNA coding and intergenic region. We additionally considered mutations within 100-bp upstream of the transcription start site of each operon, which are expected to contain promoter and regulatory sequences. The *M. tuberculosis* H37Rv annotated genome (release 4, 2021-03-23) was downloaded from https://mycobrowser.epfl.ch/releases and used as a source of annotated CDS. The transcriptional units (operons) of *M. tuberculosis* H37Rv were downloaded from https://mycobacterium.biocyc.org/. We tested CDS and operons for an excess of protein-altering (functional) mutations compared to the rest of the genome, considering the length of CDS, or cumulative length of CDS if testing operons involving multiple CDS. To do this, we performed a single-tailed Poisson test using the genome-wide mutation count per bp multiplied by the gene length as the expected number of mutations as previously implemented. P values were corrected for multiple testing using a Benjamini & Hochberg correction using the total number of functional units in the genome as the number of tests. We chose a significance level of 0.05 and reported hits with an adjusted P value below this value, unless otherwise stated.

### Determination of drug-resistance acquisition rates

Anti-TB treatment data were extracted from either TB Portals or published studies when available (Supplementary Table 3), and considered all drugs dispensed to each patient over the course of their TB treatment. Binary variables were created for each drug to indicate whether each patient had received that drug or not. The variable for fluoroquinolones was recoded so that it included levofloxacin, moxifloxacin, and ofloxacin. The variable for protionamide was also combined into one variable with ethionamide. To determine whether resistance acquisition rates, a frequency table was generated. All genes known to be associated with the emergence of DR were categorized according to the drug to which they conferred resistance to, based on existing literature (Supplementary Table 2). The number of strains that had a fixed mutation on any of the genes known to be associated with resistance to each drug were then tallied. This process was done for amikacin, bedaquiline, capreomycin, clofazimine, cycloserine, ethambutol, ethionamide, isoniazid, kanamycin, fluoroquinolones, linezolid, pyrazinamide, rifampicin, and streptomycin. The proportion was calculated as the number of strains treated with a drug and had a fixed *de novo* mutation on any gene conferring resistance to that drug divided by the number of strains treated with that drug. For each proportion, a 95% confidence interval (95% CI) was calculated.

### Source of protein structures

The crystal structure of *M. tuberculosis* WhiB6 and σA4 complex was obtained from the PDB (PDB 8DV5). AlphaFold was used to generate 3D protein models for *M. tuberculosis* EccE_1_, EspK and MycP_1_. The AlphaFold model of *M. tuberculosis* MycP_1_ showed high structural conservation with the *M. smegmatis* orthologue (PDB 4KPG, Solomonson et al., 2013) with an r.m.s.d. of 0.299 Å over 312 Cα atoms, supporting the use of this model to map the mutations identified in this study. The structural model of the *M. tuberculosis* PhoR homodimer was predicted by AlphaFold. The dimerization-phosphorylation domain of PhoR was obtained from its crystal structure (PDB 5UKV).

## Acknowledgements

This publication presents independent research supported in part by a Wellcome grant 204928/Z/16/Z awarded to Francesc Coll. The views expressed in this publication are those of the author(s) and not necessarily those of the funders. Data were obtained from the TB Portals (https://tbportals.niaid.nih.gov), which is an open-access TB data resource supported by the National Institute of Allergy and Infectious Diseases (NIAID) Office of Cyber Infrastructure and Computational Biology (OCICB) in Bethesda, MD. These data were collected and submitted by members of the TB Portals Consortium (https://tbportals.niaid.nih.gov/Partners). Investigators and other data contributors that originally submitted the data to the TB Portals did not participate in the design or analysis of this study.

## Data availability statement

The whole genome sequences of the isolate collections used in this study are available on European Nucleotide Archive (ENA) under the accessions listed in Supplementary Data 1, which also includes isolate metadata. All scripts necessary to run the described analyses are available on GitHub (https://github.com/francesccoll/mtbc-adaptive-mutations/). The list of raw, filtered, and drug resistance mutations identified to have been originated *de novo* during infection can be found in Supplementary Data 2. The full list of genes, operons and promoter regions enriched by mutations can be found in Supplementary Data 3. Supplementary Data 4 includes drug treatment data for individual strains.

## Author Contributions

Conceptualization: FC; Data curation: FC, HZ and NMJ; Formal bioinformatic analysis: FC, HZ and NMJ; Funding acquisition: FC; Investigation: FC, HZ, NMJ and AFN; Bioinformatics methodology: FC, HZ and NMJ; Laboratory methodology: not applicable; Project administration: not applicable; Resources: FC and EMH; Supervision: FC; Validation: not applicable; Visualization: HZ, NMJ and AFN; Writing – original draft: FC; Writing – review & editing: all authors.

