## Supplementary Materials for "*In host* mutational adaptation of *Mycobacterium tuberculosis* complex strains"

### Supplementary Tables

Supplementary Table 1. Number of MTBC isolate genomes and TB patients available per study before and after genomic QC.

Supplementary Table 2 *M. tuberculosis* genes known to confer resistance to anti-tuberculosis drugs.

Supplementary Table 3. Data sources and treatment data availability.

### Supplementary Figures

Supplementary Figure 1 Visual representation of genomic QC metrics and thresholds.

Supplementary Figure 2. Descriptive characteristics of all strains with available drug treatment data.

### Supplementary files

Supplementary Data 1. List of isolate genomes, accessions and metadata used in this study.

Supplementary Data 2. List of raw, filtered, and drug resistance mutations identified to have been originated *de novo* during infection.

Supplementary Data 3. List of genes, operons and promoter regions enriched by mutations.

Supplementary Data 4. Drug treatment data.

### Supplementary References

**Supplementary Table 1.** Number of MTBC isolate genomes and TB patients available per study before and after genomic QC

| | Before genomic QC | | | | After genomic QC | | | | | |
|---|---|---|---|---|---|---|---|---|---|---|
| | N isolates | | | N hosts | N isolates | | | | | N hosts |
| Study ID (citation) | total | context | within host | total | removed | total | context | same strain | different strain | kept |
| TB portals[1] | 3,211 | 2,355 | 856 | 383 | 104 | 3,107 | 2,298 | 630 | 179 | 276 |
| cox2021[2] | 1,410 | 1,174 | 236 | 118 | 40 | 1,370 | 1,146 | 192 | 32 | 96 |
| nimmo2020[3] | 615 | 246 | 369 | 118 | 107 | 508 | 229 | 277 | 2 | 95 |
| wollenberg2017[4] | 447 | 0 | 447 | 97 | 26 | 421 | 1 | 409 | 11 | 95 |
| walker2013[5] | 388 | 166 | 222 | 86 | 8 | 380 | 159 | 210 | 11 | 81 |
| walker2018b[6] | 1,555 | 1,338 | 217 | 89 | 54 | 1,501 | 1,292 | 200 | 9 | 81 |
| lieberman2016[7] | 2,693 | 0 | 2,693 | 44 | 214 | 2,479 | 0 | 2,478 | 1 | 44 |
| guerra2014[8] | 168 | 0 | 168 | 84 | 39 | 129 | 31 | 64 | 34 | 32 |
| chen2021[9] | 83 | 0 | 83 | 31 | 13 | 70 | 3 | 66 | 1 | 28 |
| perez-lago2021[10] | 68 | 0 | 68 | 27 | 5 | 63 | 2 | 61 | 0 | 25 |
| dippenaar2019[11] | 50 | 0 | 50 | 25 | 3 | 47 | 3 | 42 | 2 | 21 |
| asare2021[12] | 60 | 0 | 60 | 29 | 0 | 60 | 0 | 42 | 18 | 20 |
| casali2016[13] | 38 | 0 | 38 | 19 | 0 | 38 | 0 | 38 | 0 | 19 |
| witney2017[14] | 69 | 1 | 68 | 34 | 21 | 48 | 12 | 36 | 0 | 18 |
| liu2020[15] | 804 | 0 | 804 | 18 | 40 | 764 | 0 | 764 | 0 | 18 |
| folkvardsen2020[16] | 82 | 0 | 82 | 32 | 29 | 53 | 11 | 38 | 4 | 17 |
| shanmugam2021[17] | 82 | 0 | 82 | 41 | 25 | 57 | 17 | 28 | 12 | 14 |
| chen2020[18] | 151 | 102 | 49 | 21 | 20 | 131 | 90 | 34 | 7 | 14 |
| ssengooba2016b[19] | 26 | 0 | 26 | 13 | 2 | 24 | 2 | 22 | 0 | 11 |
| hu2021[20] | 130 | 0 | 130 | 63 | 31 | 99 | 19 | 22 | 58 | 11 |
| clark2013[21] | 51 | 36 | 15 | 5 | 0 | 51 | 36 | 13 | 2 | 4 |
| mokrousov2020[22] | 20 | 0 | 20 | 6 | 4 | 16 | 2 | 14 | 0 | 4 |
| acosta2022[23] | 33 | 1 | 32 | 3 | 3 | 30 | 1 | 29 | 0 | 3 |
| seraphin2019[24] | 31 | 0 | 31 | 3 | 0 | 31 | 0 | 31 | 0 | 3 |

| | | | | | | | | | | |
|---|---|---|---|---|---|---|---|---|---|---|
| xu2018[25] | 18 | 0 | 18 | 4 | 3 | 15 | 0 | 12 | 3 | 3 |
| mehaffy2014[26] | 66 | 60 | 6 | 3 | 19 | 47 | 43 | 4 | 0 | 2 |
| merker2013[27] | 13 | 0 | 13 | 4 | 3 | 10 | 1 | 7 | 2 | 2 |
| wada2015[28] | 10 | 8 | 2 | 1 | 0 | 10 | 8 | 2 | 0 | 1 |
| cancino2019[29] | 16 | 0 | 16 | 1 | 0 | 16 | 0 | 15 | 1 | 1 |
| eldholm2014[30] | 18 | 0 | 18 | 1 | 0 | 18 | 0 | 18 | 0 | 1 |
| ley2021[31] | 12 | 10 | 2 | 1 | 1 | 11 | 9 | 2 | 0 | 1 |
| black2015[32] | 4 | 0 | 4 | 2 | 1 | 3 | 1 | 2 | 0 | 1 |
| hjort2021[33] | 77 | 0 | 77 | 1 | 1 | 76 | 0 | 76 | 0 | 1 |
| martin2018[34] | 25 | 21 | 4 | 1 | 20 | 5 | 1 | 4 | 0 | 1 |
| liu2015[35] | 5 | 0 | 5 | 1 | 5 | 0 | 0 | 0 | 0 | 0 |
| Total | 12,529 | 5,518 | 7,011 | 1,409 | 841 | 11,688 | 5,417 | 5,882 | 389 | 1,044 |

The “within host” isolate genomes are those taken from TB patients who had multiple isolates sequenced per host. The total number of these patients per study is indicated in the fifth column (N hosts total). In addition, isolate genomes from other patients who had a single MTBC genome sequenced per host were considered as contextual isolates (“context”). The total number of isolates before genomic QC corresponds to the sum of “within host” and “context” isolates. After genomic QC, the genomes that did not meet the genomic QC criteria were discarded (“removed”). The “within host” isolate genomes were split between those assigned to a different strain (“different strain”) and those assigned to the same strain (“same strain”), which were kept for further analyses. The number of patients with multiple isolates sequenced of the same strain is indicated in “N hosts kept” (total 1,044 patients across studies).

**Supplementary Table 2** *M. tuberculosis* genes known to confer resistance to anti-tuberculosis drugs

| Drug | Line | Locus tag (gene name)* |
|---|---|---|
| Amikacin | 2nd | Rv0529 (*ccsA*), **Rv2416c (*eis*) promoter**, Rv3106 (*fprA*), **MTB000019 (*rrs*)**, Rv3805c (*aftB*), Rv3862c (*whiB6*), Rv3197A (*whiB7*) |
| Bedaquiline | last | Rv1305 (*atpE*), Rv2535c (*pepQ*), Rv1979c, **Rv0678**, Rv0676c (*mmpL5*), Rv0677c (*mmpS5*) |
| Capreomycin | 2nd | Rv0529 (*ccsA*), Rv3106 (*fprA*), **MTB000019 (*rrs*)**, **Rv1694 (*tlyA*)**, Rv3805c (*aftB*), Rv3862c (*whiB6*) |
| Clofazimine | last | Rv2535c (*pepQ*), Rv1979c, **Rv0678**, Rv0676c (*mmpL5*), Rv0677 (*mmpS5*) |
| Cycloserine | last | **Rv2780 (*ald*)**, **Rv3423c (*alr*)** |
| Delamanid | last | Rv3547 (ddn), Rv3261- Rv3262 (fbiA-fbiB), Rv1173 (fbiC), Rv2983, Rv0407 (fgd1) |
| Ethambutol | 1st | **Rv3795 (*embB*)**, Rv3793 (*embC*), Rv3794 (*embA*), Rv3792 (*aftA*), Rv1267c (*embR*), Rv3806c (*ubiA*) |
| Ethionamide | 2nd | Rv3083, Rv1484 (*inhA*), Rv0486 (*mshA*), Rv1854c (*ndh*), **Rv3854c (*ethA*)**, **Rv3854c (*ethA*) promoter,** Rv3855 (*ethR*), Rv0565c |
| Isoniazid | 1st | **Rv1908 (*katG*)**, Rv2428 (*aphC*), **Rv2428 (*aphC*) promoter**, Rv1484 (*inhA*), **Rv1484 (*inhA*) promoter**, Rv0486 (*mshA*), Rv1854c (*ndh*), Rv1258c, Rv2752c |
| Kanamycin | 2nd | **MTB000019 (*rrs*)**, **Rv2416c (*eis*) promoter**, Rv3197A (*whiB7*) |
| Fluoroquinolones | 2nd | **Rv0006 (*gyrA*), Rv0005 (*gyrB*)** |
| Linezolid | last | Rv0701 (*rplC*), MTB000020 (*rrl*) |
| PAS** | last | **Rv2764c (*thyA*)[36], Rv2754c (*thyX*)[37] promoter** |
| Pyrazinamide | 1st | Rv3596c (*clpC1*), Rv3601 (*panD*), **Rv2043c (*pncA*)**, Rv1918c (*PPE35*), Rv1258c, Rv3236c |
| Rifampicin | 1st | **Rv0667 (*rpoB*)**, **Rv0957 (*rpoC*)**, Rv3457c (*rpoA*), Rv2752c |
| Streptomycin | 2nd | **Rv3919c (*gid*)**, **Rv0682 (*rpsL*)**, **MTB000019 (*rrs*),** Rv1258c Rv3862c (*whiB6*), Rv3197A (*whiB7*) |

* Whole list of genes extracted from. In bold, drug resistance genes detected in this study with signals of *in host* mutational adaption. Genes approaching statistical significance in this analysis are underscored. **PAS, para-aminosalicylic acid. Genes linked to PAS resistance not included in[1] but identified from other sources.[36,37]

**Supplementary Table 3.** Data sources and treatment data availability.

| Study ID (citation) | Number of strains | Treatment data |
|---|---|---|
| acosta2022[23] | 3 | *Not available* |
| asare2021[12] | 20 | Available |
| black2015[32] | 1 | *Not available* |
| cancino2019[29] | 1 | Available |
| casali2016[13] | 19 | *Not available* |
| chen2020[18] | 14 | *Not available* |
| chen2021[9] | 28 | *Not available* |
| clark2013[21] | 5 | Available |
| cox2021[2] | 96 | *Not available* |
| dippenaar2019[11] | 21 | *Not available* |
| eldholm2014[30] | 1 | *Not available* |
| folkvardsen2020[16] | 12 | *Not available* |
| guerra2014[8] | 32 | Available |
| hjort2021[33] | 1 | Available |
| hu2021[20] | 11 | Available |
| ley2021[31] | 1 | *Not available* |
| lieberman2016[7] | 45 | *Not available* |
| liu2020[15] | 19 | *Not available* |
| martin2018[34] | 1 | *Not available* |
| mehaffy2014[26] | 12 | *Not available* |
| merker2013[27] | 2 | *Not available* |
| mokrousov2020[22] | 4 | Available |
| nimmo2020[3] | 95 | Available |
| perez-lago2021[10] | 25 | *Not available* |
| seraphin2019[24] | 3 | *Not available* |
| shanmugam2021[17] | 14 | *Not available* |
| ssengooba2016b[19] | 11 | *Not available* |
| TB Portals[1] | 285 | Available |
| wada2015[28] | 1 | *Not available* |
| walker2013[5] | 81 | *Not available* |
| walker2018b[6] | 81 | *Not available* |
| witney2017[14] | 18 | Available |
| wollenberg2017[4] | 99 | Available |
| xu2018[25] | 3 | Available |

**Supplementary Figure 1** Visual representation of genomic QC metrics and thresholds

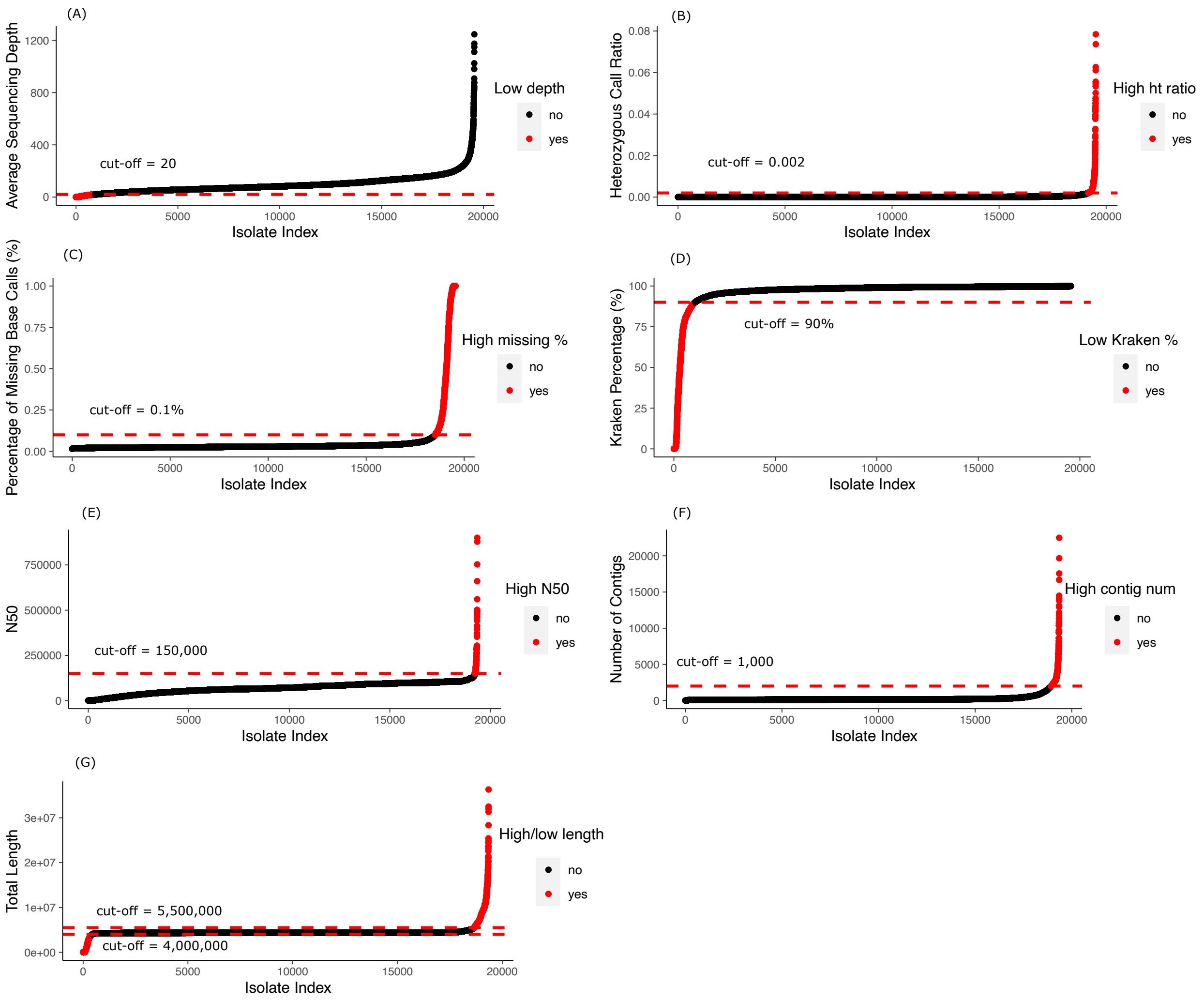

The scatter plots of individual genomic QC metrics are shown as different panels: (A) Average sequencing depth, (B) heterozygous call ratio, (C) missing base call percentage, (D) Kraken percentage of MTBC reads, (E) assembly N50, (F) number of contigs in assembly, and (G) total length of assembled genome. The values for each QC metric were plotted in a scatter plot to visually identify outliers. In most cases, a cut-off line can be drawn to easily separate outliers from the rest and set QC thresholds. For average sequencing depth, a cut-off of 20 was used in this study.

**Supplementary Figure 2.** Descriptive characteristics of all strains with available drug treatment data.

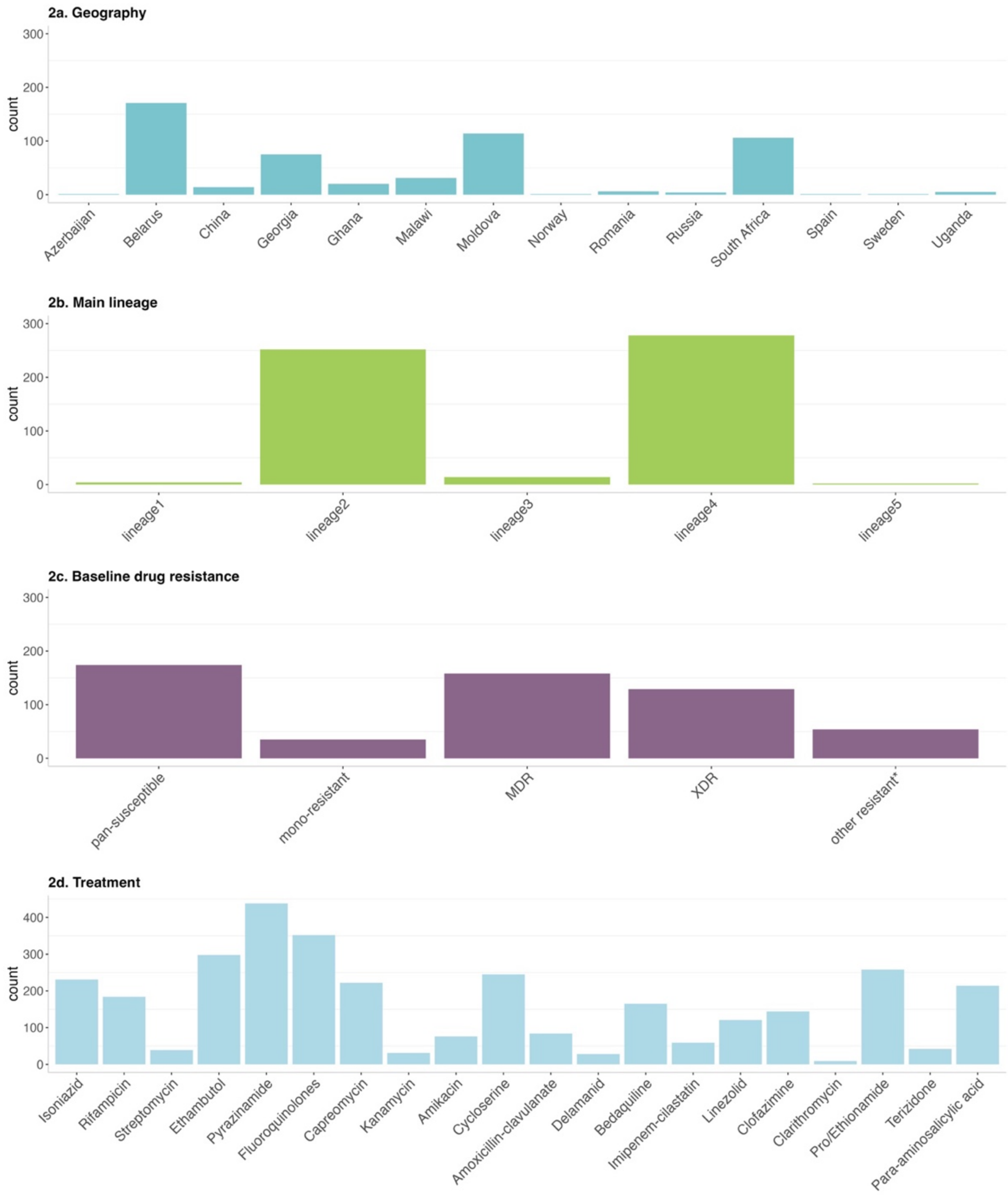

**Other resistant* refers to any strains that had predicted resistance to more than one drug but did not meet the clinical MDR or XDR definitions.
